# Engineered cell-to-cell signalling within growing bacterial cellulose pellicles

**DOI:** 10.1101/372797

**Authors:** Kenneth T Walker, Vivianne J Goosens, Akashaditya Das, Alicia E Graham, Tom Ellis

**Author notes:** These authors contributed equally to this work (Kenneth T Walker, Vivianne J Goosens).

## Abstract

Bacterial cellulose is a strong and flexible biomaterial produced at high yields by *Acetobacter* species and has applications in healthcare, biotechnology and electronics. Naturally, bacterial cellulose grows as a large unstructured polymer network around the bacteria that produce it, and tools to enable these bacteria to respond to different locations are required to grow more complex structured materials. Here, we introduce engineered cell-to-cell communication into a bacterial cellulose-producing strain of *Komagataeibacter rhaeticus* to enable different cells to detect their proximity within growing material and trigger differential gene expression in response. Using synthetic biology tools, we engineer Sender and Receiver strains of *K. rhaeticus* to produce and respond to the diffusible signalling molecule, acyl-homoserine lactone (AHL). We demonstrate that communication can occur both within and between growing pellicles and use this in a boundary detection experiment, where spliced and joined pellicles sense and reveal their original boundary. This work sets the basis for synthetic cell-to-cell communication within bacterial cellulose and is an important step forward for pattern formation within engineered living materials.

## Introduction

Cellulose is a simple, yet versatile glucose polymer from which biology weaves a broad array of materials. It is one of nature’s most abundant polymers, and can form the basis of materials that are light and elastic, such as a loofah, or structures that are strong and stiff like bamboo (Youssefian *et al*., 2015; Chen *et al*., 2017). Cellulose is produced most abundantly by plants, however, a number of bacterial species naturally overproduce cellulose as part of their biofilm matrix, making a structure known as a pellicle (Römling *et al*., 2015). Bacterial cellulose differentiates itself from plant cellulose in that it is typically produced as ultrapure nanocellulose fibres, free from the contaminating polymers like pectin and lignin that are co-produced by plant cells. Bacterial cellulose (BC) is well known for being strong and flexible, with a single nanofiber having the tensile strength of ~1 GPa and a Young’s modulus of 114 GPa; whilst also being highly hydrophilic, with water making ~90% of its weight in its wet state (Lee *et al*., 2014). These material properties have seen commercial uptake of bacterial cellulose for biomedical and cosmetic applications, such as for protective bandaging, while also offering potential as a material used in electronics, e.g. as a battery separator or as a matrix ingredient in organic light-emitting diode (OLED) displays (Lee *et al*., 2014; Jang *et al*., 2017). Some BC-producing bacteria have also been demonstrated to be amenable to genetic engineering (Florea *et al*., 2016), offering the possibility of using cellulose-producing bacteria as a means to produce genetically-defined materials, where the material properties and functions can be programmed by engineering at the DNA level using the modern tools of synthetic biology (Cameron *et al*., 2014). Programming material properties and features into biomaterials using such tools is part of the emerging new field of engineered living materials (ELMs) (Nguyen *et al*., 2018).

In most natural systems that produce biomaterials, communication between material-producing cells is a critical component of producing materials with structures on the micro and macro scale. ELMs research can theoretically recapitulate this in material-producing bacteria by leveraging the significant past work done to engineer *Escherichia coli* for synthetic pattern formation (Scholes *et al*., 2017). Such engineered pattern formation work typically exploits quorum-sensing systems from various bacteria, which are a natural method of bacterial cell-to-cell communication that can be reprogrammed (Scholes *et al*., 2017). The Lux quorum-sensing system from *Vibrio fischeri* is particularly well-used in this context, utilising the signalling molecule 3OC6-HSL – often referred to generally as acyl-homoserine lactone (AHL) (Churchill *et al*., 2011). AHL is produced by the enzyme LuxI, this diffusible molecule can then bind and activate the transcription factor LuxR, which in activates the pLux promoter to drive expression of a downstream gene (Waters *et al*., 2005). The diffusible and inducible nature of AHLs can be exploited to create a morphogen gradient - the basis for many models of patterning, such as the French Flag Model (Wolpert, 1969) and Turing patterns (Turing, 1952) (Fig. 1A). In synthetic pattern formation research, the Lux system has been used to produce programmable bullseye patterns on dishes of engineered *E. coli* cells that produce AHL (‘Senders’), while being surrounded by a lawn of cells that express fluorescent proteins in response to different levels of AHL (‘Receivers’) (Basu *et al*., 2005). Since then, more complex systems have been engineered using AHL signalling in *E. coli*, including tuneable stripe patterns with motile cells (Liu *et al*., 2011), hierarchical patterning with multiple AHLs (Boehm *et al*., 2018) and the recent creation of stochastic Turing patterns (Karig *et al*., 2018).

**Figure 1.**
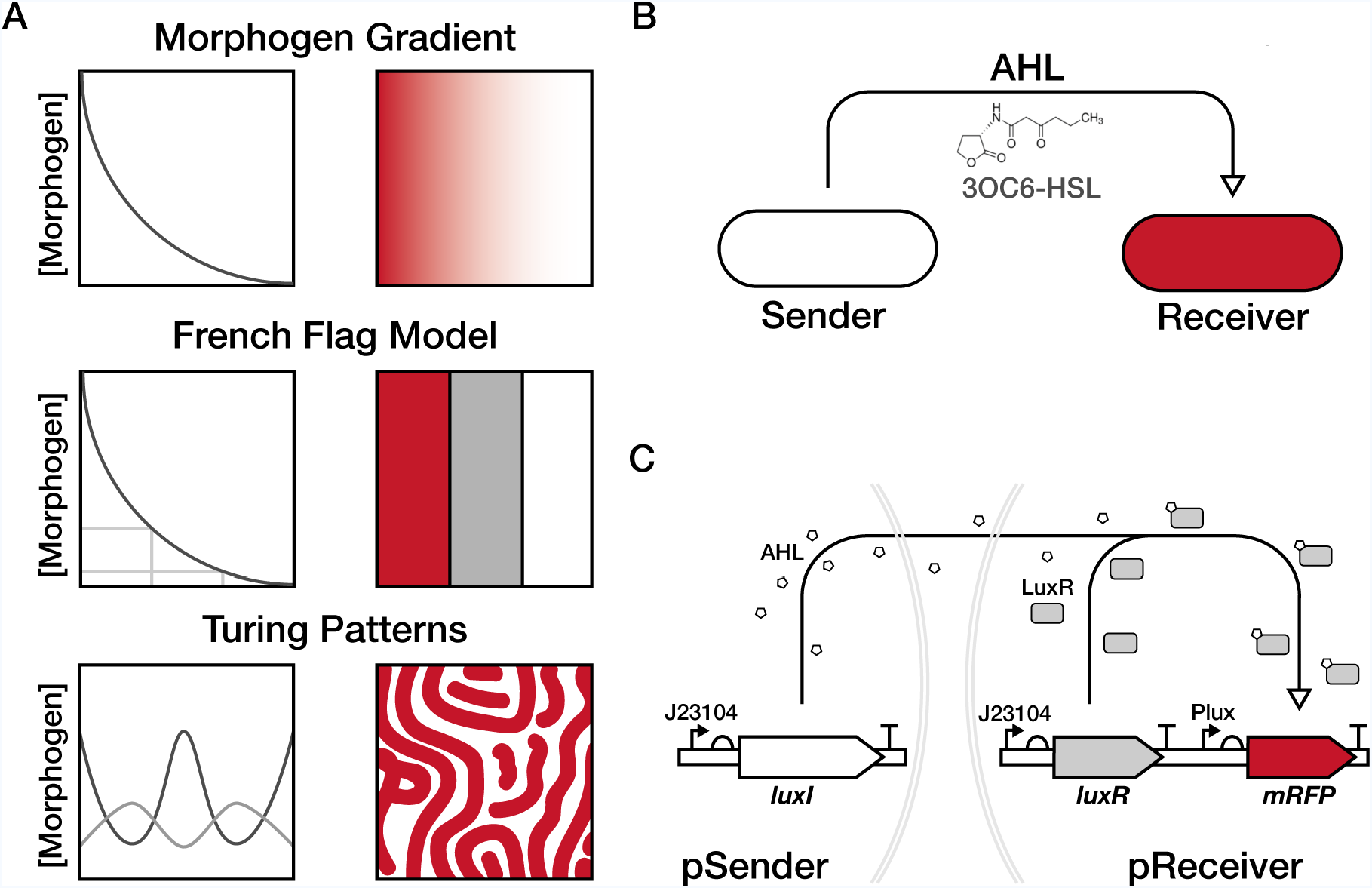
Design of a unidirectional cell-to-cell communication system. **(A)** Illustration of diffusion driven patterning models. At the top is a morphogen gradient, a simple form of patterning that is also the basis for the French Flag model, presented immediately below, and the even more complex Turing patterns presented in the bottom row. The French Flag model requires sensing the concentration of a morphogen to differentiate the response, whilst Turing patterns require two morphogens that regulate each other and diffuse at different rates. **(B)** The cell-to-cell communication system use here is made up of two different cells, a Sender cell producing the N-(3-oxohexanoyl) homoserine lactone AHL molecule and a Receiver cell that activates *mRFP* expression (red) in response to that AHL. **(C)** The individual plasmid constructs used in this study. The pSender plasmid constitutively expresses the AHL synthase LuxI, and the pReceiver plasmid constitutively expresses the transcriptional activator LuxR. When LuxR is bound to AHL it up regulates expression from the PLux promoter which is this case activates expression of a monomeric red fluorescent protein (mRFP) gene. Both *luxI* and *luxR* are expressed in these constructs with constitutive promoter J23104.

Previously we isolated a genetically-tractable BC-producing *Komagataeibacter rhaeticus* strain and developed a basic kit of synthetic biology parts and tools to enable its genetic engineering (Florea *et al*., 2016). Here, we extend our initial *K. rhaeticus* toolkit to now add engineered cell-to-cell communication to enable downstream work in patterning structural and functional properties into BC-based ELMs. By modifying the AHL-based systems developed previously for *E. coli*, we introduce and characterise cell-to-cell communication between engineered sender and receiver *K. rhaeticus* cells and demonstrate this system to be active within pellicles and able to define the boundaries between pellicle regions.

## Results

### Construction of Sender and Receiver plasmids

To establish synthetic cell-to-cell communication in *K. rhaeticus*, we designed two separate strains, a Sender strain that produces the diffusible AHL molecule N-(3-oxohexanoyl) homoserine lactone (3OC6-HSL), and a Receiver strain that can sense AHL and respond by expressing an easy-to-measure reporter gene (Fig. 1B). The sender plasmid, pSender, was constructed by placing the AHL-synthesis gene *luxI* downstream of the synthetic constitutive promoter J23104 (Fig. 1C). The receiver plasmid, pReceiver, used promoter J23104 for expression of the gene *luxR* and the LuxR-regulated promoter, pLux for expression of the monomeric red fluorescent protein (*mRFP*) gene (Fig. 1C). The J23104 constitutive promoter was used in both instances as it was previously shown to have high activity in *K. rhaeticus* (Florea *et al*., 2016).

### AHL production by Sender strains

We first determined whether our engineered Sender *K. rhaeticus* cells produced and released meaningful quantities of AHL to trigger cell-to-cell communication. To measure effective AHL production levels we used *E. coli* transformed with pReceiver as an assay system, as *E. coli* with similar receiver plasmids give a well-characterised increase in reporter gene expression in response to increasing amounts of AHL (Canton *et al*., 2008). Standard concentrations of chemically-produced AHL were added to the growth media of *E. coli* Receiver to produce a dose response curve where relative mRFP production levels in *E. coli* correspond to known levels of AHL (Fig. 2A). We next grew *K. rhaeticus* Sender cells, and a control *K. rhaeticus* containing only an empty plasmid (pEmpty), collected their spent growth media and used the *E. coli* Receiver cells to estimate the AHL concentrations produced. For the *K. rhaeticus* Sender cells, spent growth media induced mRFP production in *E. coli* that matched nanomolar levels of synthetic AHL on the dose response curve (Fig. 2A.). Interestingly, detectable levels of *E. coli* mRFP production were also observed when spent growth media from the *K. rhaeticus* pEmpty control cells were used, but not when unused HS media was added.

**Figure 2.**
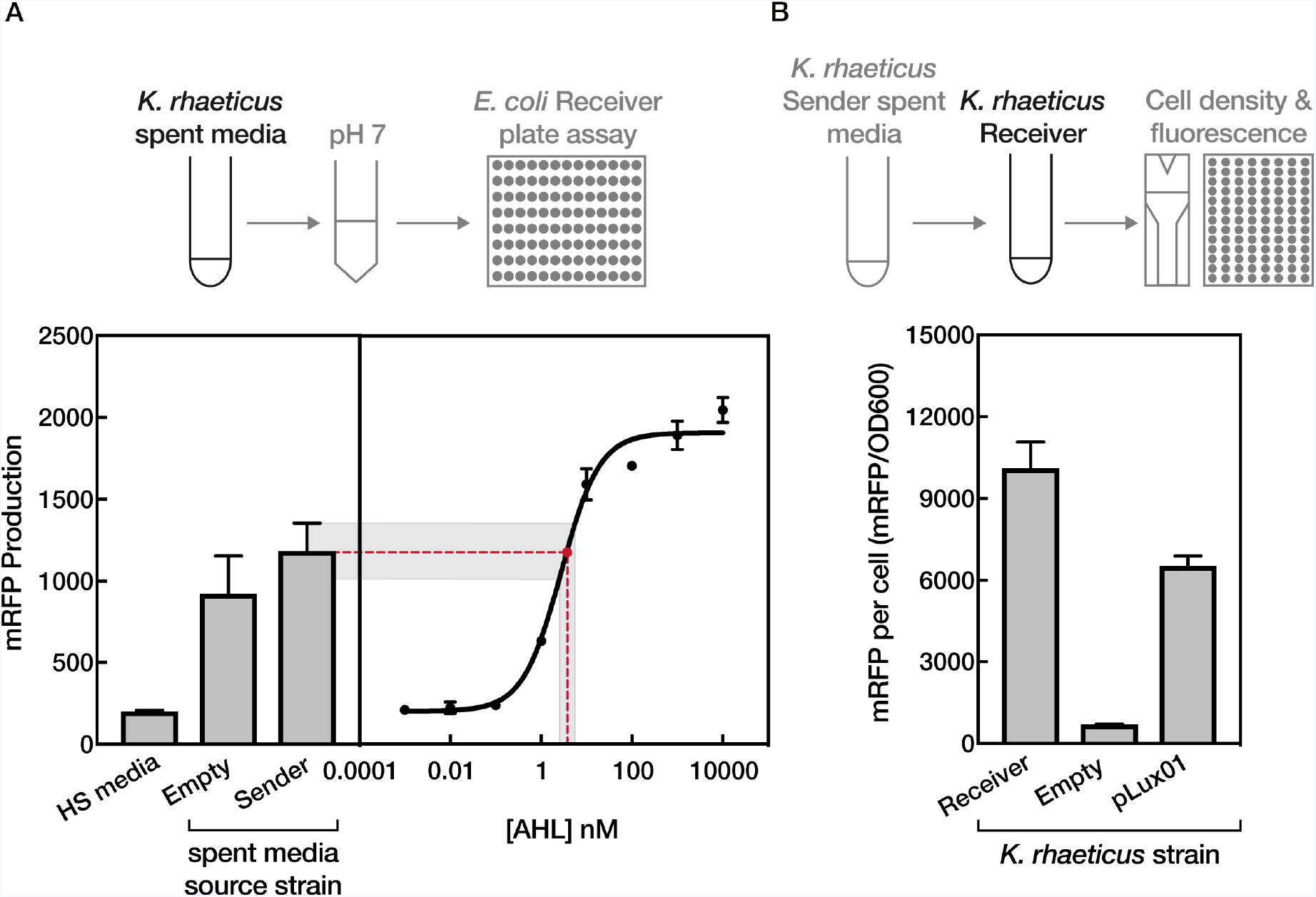
Production and effect of AHL produced by *K. rhaeticus* Sender strains. **(A)** Testing the ability of *K. rhaeticus* Sender to produce AHL. The top illustration demonstrates the process of testing the AHL concentration in *K. rhaeticus* spent media. The left chart shows the mean mRFP fluorescence response from an *E. coli* Receiver assay after the addition of fresh HS media and spent media from *K. rhaeticus* Empty and *K. rhaeticus* Sender. The dose response curve is based on the *E. coli* response to known concentrations of added AHL standards and is used to estimate the approximate concentration of AHL produced by *K. rhaeticus* Sender. **(B)** The mean mRFP production of liquid *K. rhaeticus* Receiver strains upon addition of spent media from *K. rhaeticus* Sender. The top illustration details the process of adding spent media to *K. rhaeticus* and the methods used to measure the fluorescence response. Treated strains included *K. rhaeticus* Receiver, the original pLux01 vector and *K. rhaeticus* with pEmpty. Error bars denote the standard deviation from three repeats.

Next, the engineered *K. rhaeticus* Receiver cells were assessed for their ability to respond to AHL and express the mRFP reporter gene. Spent media from *K. rhaeticus* Sender was applied to *K. rhaeticus* Receiver liquid cultures (and control cells) and mRFP production levels determined. *K. rhaeticus* Receiver strains produced significant levels of mRFP in response to *K. rhaeticus* Sender spent media, giving approximately 10-fold more red fluorescence than background levels (by comparing pEmpty control strains against *K. rhaeticus* Receiver). The engineered pReceiver plasmids, using the J23104 promoter for *luxR* expression, also showed improvement in the sensitivity of the AHL-response system in *K. rhaeticus*, producing more mRFP from the same input when compared to the previously described pLux01 construct (Florea *et al*., 2016) that uses a weaker promoter for *luxR* expression (Fig. 2B).

### Cell-to-cell communication between Sender and Receiver strains

Having determined that Sender and Receiver *K. rhaeticus* could produce and respond to appropriate levels of AHL in isolation, we next examined direct cell-to-cell communication between mixed cultures of the two strains. To investigate interactions at the single cell level, we first examined our co-cultures via fluorescence microscopy. This was enabled using a microfluidic plate set-up that provides continual slow perfusion of new growth media. The set-up allowed *K. rhaeticus* to grow in a single focal plane and yielded single-cell resolution images of the cultures. Sender and Receiver *K. rhaeticus* strains were mixed at low density in a 1:1 ratio and incubated at 30 °C in the microfluidic plate system for 48 hours. After this, images of the co-culture were taken and these revealed strong expression of mRFP in clusters of cells surrounded by those not showing any red signal (Fig. 3A). These expressing clusters are presumably Receiver cells that have grown from an ancestor while surrounded by the non-fluorescent Sender cells. To verify this, we imaged Receiver cells grown mixed with pEmpty control cells and exposed them to 100 nM of synthetic AHL and saw the same pattern. As a further control, cells of only the *K. rhaeticus* Receiver strain were also grown in the set-up and showed only very low-level background red fluorescence (Fig. 3A), confirming that only engineered expression of AHL from Sender strains can trigger significant expression of the reporter gene in Receiver *K. rhaeticus*.

**Figure 3.**
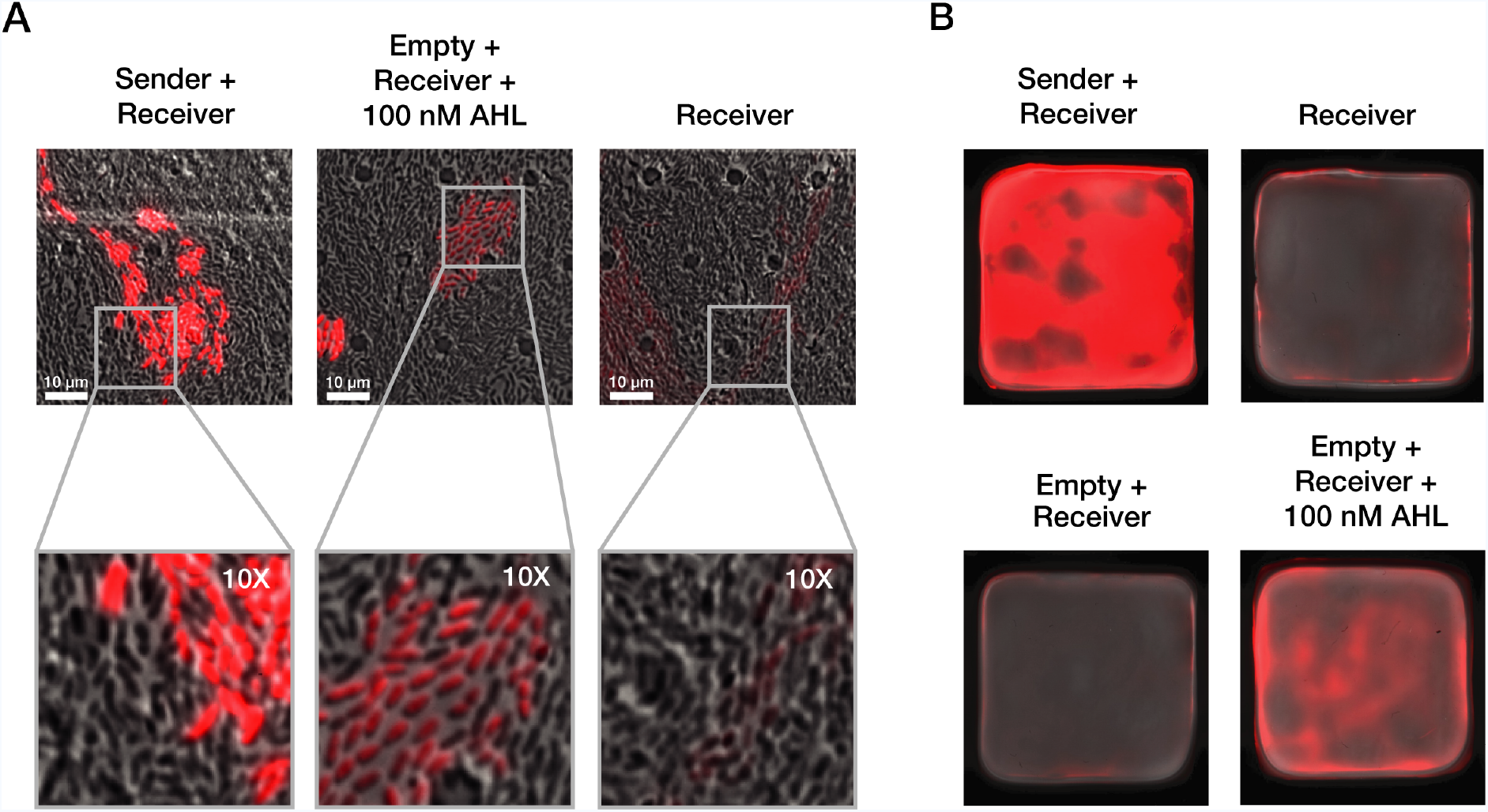
Cell-to-cell signalling functions within co-cultured pellicles. **(A)** Fluorescence microscopy of co-cultures. Images detail *K. rhaeticus* growth after 48 hours within three separate chambers on the same microfluidics plate. Images were taken with a 60X oil emulsion lens. The bottom column displays a digitally zoomed in region of the images in the top. All microscopy images were taken with the same settings for brightfield and mRFP fluorescence channels. Brightness and contrast for the brightfield channel were adjusted to improve clarity, while the red fluorescence channel was left unadjusted **(B)** Pellicle co-cultures. The top left pellicle is a 50:50 mix of Sender and Receiver culture and the top right a pure Receiver pellicle. Both the bottom pellicles are a 50:50 mix of Empty and Receiver, where the bottom left was left to grow without AHL while the bottom right had 100 nM AHL added to it on second day of growth.

The results so far showed that our engineered *K. rhaeticus* behave similarly to *E. coli* engineered with similar constructs in past work, however, unlike *E. coli, K. rhaeticus,* naturally produces and grows within a macro-scale material structure. Whilst giving many benefits in terms of downstream applications, the growth of the bacteria in a pellicle may alter the way that gene circuits operate, for example by having the cells switch to stationary phase or making cell conditions anaerobic and thus changing the cell state. Therefore, to determine if the engineered communication system can function in this environment, we next switched to pellicle growth experiments using co-cultures of our Sender and Receiver strains. A 50:50 inoculation ratio of Sender and Receiver *K. rhaeticus* was set-up and grown alongside equivalent growth of the same control combinations used in the microscopy experiment (Fig. 3B). After 4 days this gave pellicles with the thickness of approximately 5 mm, and when scanned using a fluorescence imager, significant mRFP expression was observed across the pellicle in the Sender+Receiver combination. No fluorescence was observed with just Receiver cells or in the control pellicle where pEmpty was present in cells mixed with the *K. rhaeticus* Receiver. Fluorescence was observed when synthetic AHL was added exogenously at 100 nM AHL to this control pellicle (Fig. 3B), although the signal was not as strong as for the engineered co-culture, presumably because local AHL concentrations are higher and continually produced when it is being produced within the material.

### Cell-to-cell communication between Sender and Receiver pellicles

Past work in *E. coli* synthetic biology has established that there are many downstream applications once cells within a co-culture can be engineered to communicate (Basu *et al*., 2005; Boehm *et al*., 2018; Karig *et al*., 2018). Given the unique situation of having now established such a system in a material-producing microbe, we decided to investigate if it could be used to mediate communication between whole pieces of grown materials. In engineered living materials this could be used, for example, to determine the edges of grown areas and trigger gene expression at section boundaries. To demonstrate this, Sender and Receiver *K. rhaeticus* pellicles were grown separately, before placed together in a plate with HS media for 24 hours. Control experiments were also done in parallel, where pellicles from cells containing pEmpty were placed with Receiver pellicles and co-incubated for 24 hours with and without exogenous synthetic AHL. As before pellicles were imaged using a fluorescent scanner, and the images of the pellicle pairs show that the Receiver pellicle only activates mRFP expression when co-incubated with the Sender pellicle or when exogenous AHL was added (Fig. 4A). The signal is strongest at the pellicle edges, representing the areas that see new growth of cells and have the maximum exposure to diffusing AHL coming from the Sender pellicles.

**Figure 4.**
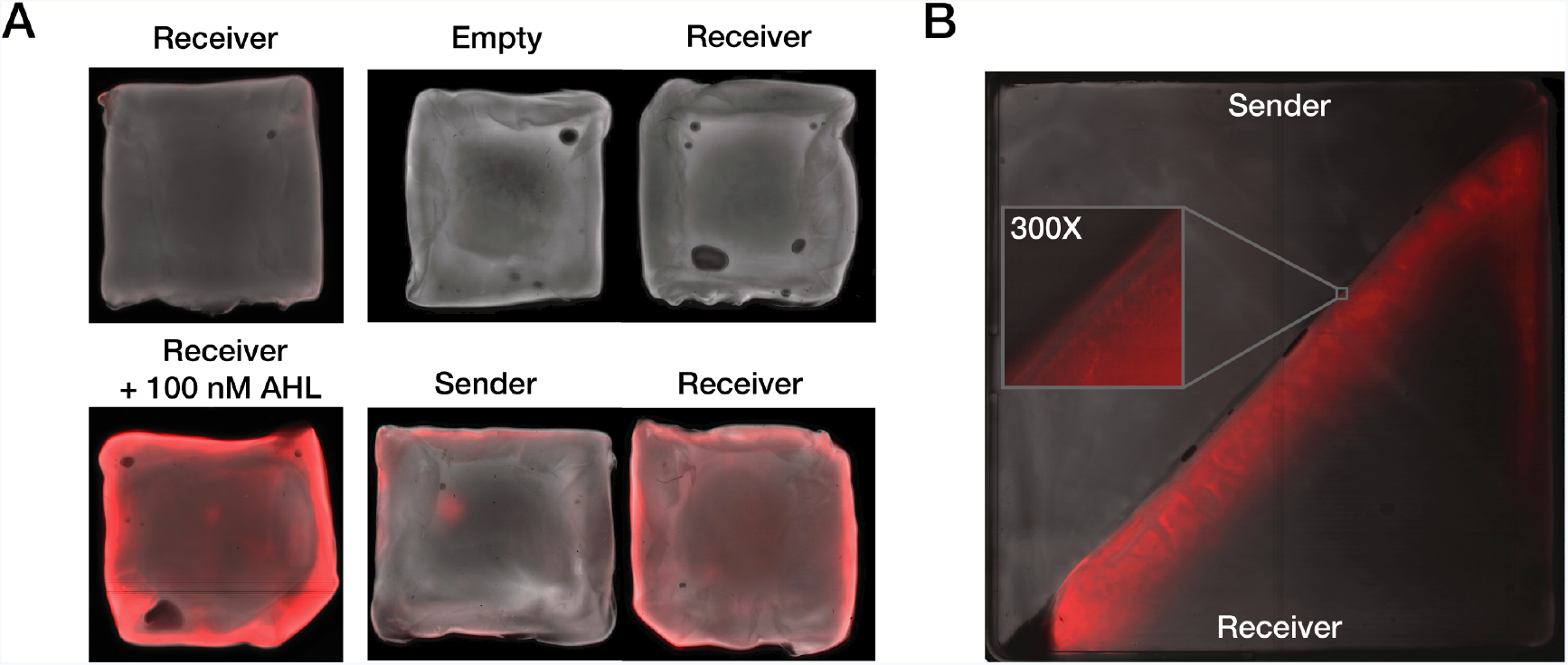
Cell-to-cell signalling can function between separate Sender and Receiver pellicles. All pellicle images have two colour channels, a pseudo-brightfield and red fluorescence channel. **(A)** Receiver pellicle induction by Sender pellicle. Left images show two Receiver pellicles after 24 hours incubation in 5 mL HS media, the top pellicle without any added AHL and the bottom pellicle with 100 nM AHL. The right images show Receiver-Sender and Empty-Sender pellicle pairs after 24 hours co-incubation in 5 ml HS media. **(B)** Boundary detection. Spliced Sender-Receiver pellicle was grown for 24 hours and then removed from soft agar surface and imaged. A digitally zoomed in region of the boundary between the two pellicles is featured.

Having verified that the Sender pellicle can produce enough AHL to diffuse out and induce gene expression in Receiver pellicles in just 24 hours, we next moved to boundary detection. In this case, cut Sender and Receiver pellicles were placed adjacent to one another and left to continue to grow in media so that they fuse together to make a larger piece. This technique of fusing pellicles via growth can aid in growing large material products. After only 24 hours incubation of closely adjacent pellicles, fusion had occurred along most of the boundary, now creating a single pellicle. This was imaged as before and revealed a clear line of strong mRFP expression along the edge of the previous Receiver pellicle. This line of expression reveals where the border of the fusion has occurred and in future engineered strains could be the location of the expression of other proteins, such as those that provide stronger material bonding (Fig. 4B).

## Discussion

Here we constructed a unidirectional cell-to-cell communication system in the cellulose producing bacteria *K. rhaeticus* using a system of two separate engineered strains: Sender and Receiver. We utilised the Lux quorum sensing system, commonly used in artificial pattern formation experiments, and showed it to be functional in *K. rhaeticus*, even within growing bacterial cellulose pellicles. Previous research on synthetic patterning of bacterial cells has entirely focused on patterns formed by colonies on agar surfaces, cells in microfluidic chambers or multi-well plates, whereas here we demonstrate this system working as the engineered cells grow in and around their own macro-material structure.

The work presented here offers a route towards the development of more intricate gene regulation and signalling circuits within growing pellicles made by *K. rhaeticus,* and will be enabled in the future by the addition of further signalling and regulatory components. Increasing the complexity of cell-to-cell communication, e.g. by bringing in a second diffusible signalling molecule, would allow the realisation of more elaborate patterns, such as Turing patterns, and also enable bidirectional communication between engineered cells. Interestingly, our initial characterisation with *E. coli* Receiver strains showed that un-engineered *K. rhaeticus* produces other molecules that partially trigger the Lux system response in *E. coli* (Fig. 2A). This hints at an a unknown native quorum system in *K. rhaeticus* which could be further investigated and potentially exploited as a second communication route. While this triggered some expression in Receiver *E. coli*, it notably did not cause a response in Receiver *K. rhaeticus* (Fig. 3), suggesting that it is not the exact same AHL molecule. A genomic search of *K. rhaeticus* only identifies an orphaned *luxR* homologue, and no obvious AHL synthases (Florea *et al*., 2016), however, recent work has shown that a closely-related *Acetobacter* strain does produce quorum molecules (Liu *et al*., 2018).

A direct potential utility of the boundary detection shown here (Fig. 4), could be to create a self-repairing material that re-joins cuts with stronger attachment than just natural pellicle fusion. This could be achieved by replacing the *mRFP* gene in the pReceiver construct by genes that express proteins that promote stronger attachment, e.g. by the surface display of adhesive catecholamines (Park *et al*., 2014) or via production of bacterial adhesive fibres such as Curli (Nguyen *et al*., 2014). Our work could also be further expanded by combining our signalling with genetic logic gates that link cell-to-cell communication with decision making and the response to combinations of external and internal stimuli (Brophy *et al*., 2014; Scott *et al*., 2016; Liu *et al*., 2018). Overall, we believe that a living bacterial cellulose material capable of programmable cell-to-cell communication could be adapted for a variety of purposes, and further engineered to create patterned materials with unique macro structure-driven material properties and novel functions.

## Experimental Procedures

### Strains, plasmids and culturing conditions

*E. coli* was cultured in at 37°C shaking at 250 rpm in Lysogeny broth (LB) (10 g/l Tryptone, 5 g/l Yeast Extract and 5 g/l NaCl) or statically at 37°C in LB agar with 34μg/ml chloramphenicol when appropriate. Cultures of *K. rhaeticus iGEM* were grown at 30°C in Hestrin – Schramm media (HS) (20g/l glucose, 10g/l yeast extract, 10g/l peptone, 2.7g/l Na2HPO4 and 1.3g/l citric acid, pH 5.6-5.8) or on HS agar plates (1,5% agar) and supplemented with 2% cellulase, when appropriate 34 μg/ml chloramphenicol was added. Electroporation of *K. rhaeticus* strains were performed as detailed previously (Florea *et al*., 2016).

For *K. rhaeticus* pellicle growth, parental pellicles were first generated by inoculating strains from glycerol stocks in 5 ml HS media and incubated at 30°C for 7 days. In order to maintain plasmids 340 ng/μl chloramphenicol was added. Liquid beneath the parent pellicle was then used, in 1:50 ratio, to inoculate parallel experimental pellicles. Experimental pellicles were harvested after 3-4 days of growth at 30°C. To generate mixed strain pellicles, the media from beneath each parent pellicle was removed and mixed at a 50:50 ratio, this mix was then inculcated into fresh HS media and placed in a deep-well 24 well plate (Axygen).

Plasmids used in this study are listed in Table 1. The vector backbones were based on previous BioBrick compatible vectors (for detailed cloning description see (Florea *et al*., 2016)) and J23104 promoter was adapted using Phusion polymerase (NEB) and PCR mutagenesis (primers listed in Table 2).

**Table 1.** Plasmids used in this study.

| Plasmids used in this study | Details | Source |
| --- | --- | --- |
| pEmpty (previously named pSEVA331Bb) | *E. coli*-*K. rhaeticus* expression vector; ori-pBRR1 origin of replication; Chloramphenicol resistant; pSEVA based plasmid with Biobrick modifications. | (Florea *et al.*, 2016) |
| pLux01 | pSEVA331Bb derivative carrying *mRFP* gene under the control of pLux and *luxR* gene under control of pLac promoter. | (Florea *et al.*, 2016) |
| pReceiver | pLux01 derivative carrying *mRFP* gene under the control of pLux and *luxR* gene under control of J23104 promoter | This study |
| pSender | pSEVA331Bb derivative carrying *LuxI* gene under the control of J23104 promoter. | This study |

**Table 2.**
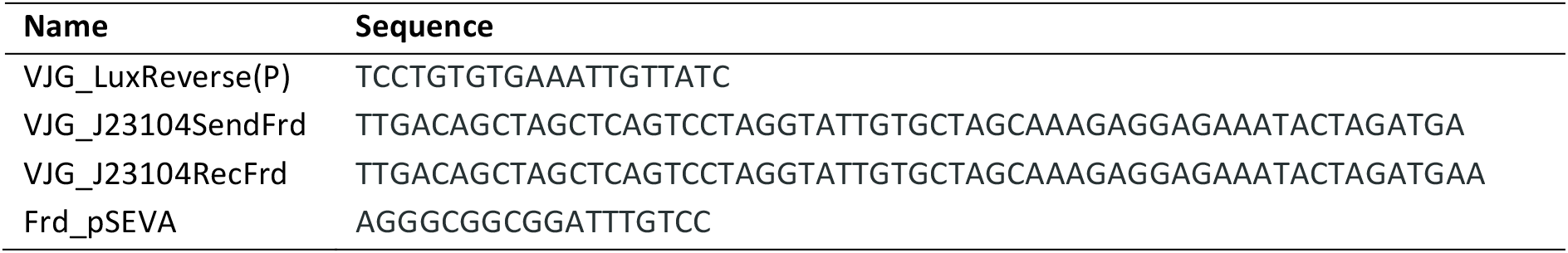
Primers used in this study.

### Spent media assays

Spent media was made by growing *K. rhaeticus* Sender strain to late exponential/early stationary phase, cells were then pelleted, and supernatant filtered before used to preform assays. Spent media or synthetic AHL (product K3255, Sigma Aldrich) was added to exponentially growing cultures and fluorescence measured after an hour. mRFP production was calculated as difference in fluorescence (measured at ex., 590 nm; em., 645 nm) between successive time points normalized by OD_600_. Multiple assays were performed in 96 well plate format in the Synergy HT Microplate Reader (BioTek) each with triplicate technical replicates. In order to estimate AHL concentration *E. coli* strain were transformed with relevant vectors, and grown in a 96 well format. Before adding to *E. coli* cultures spent HS-media was buffered to pH 7 using Tris pH 8.0, this was required as HS media has a low pH that prevent *E. coli* growth if added directly. *K. rhaeticus* strains were grown in 25 ml tubes, with vigorous shaking and cellulase, after addition of spent media samples were plated into 96 well plates and fluorescent measurements taken at ex., 590 nm; em., 645 nm within the hour. Cellulose production encouraged clumping in liquid *K. rhaeticus* cultures, making precise OD measurements for growing liquid cultures challenging, even with high cellulase treatment. Therefore, when adding spent media to *K. rhaeticus*, the mRFP response was measured within the first doubling time before clumping caused an effect.

### Fluorescence imaging

Fluorescence images of pellicles were taken with a Fujifilm FLA-5000 Fluorescent Image Analyser, with excitation at 532 nm, 600 v and Cy5 filter, whilst Pseudo-brightfeild images were taken with excitation at 473 nm, 400 v and FITC filter. Fluorescence microscopy images were taken on a Nikon Eclipse Ti inverted microscope using a Rolera EM-C^2^ camera. *K. rhaeticus* strains were grown within a ONIX microfluidics plate (B04A) (Merck) with flowing HS media at 1 psi and incubated at 30°C. brightfield and mRFP fluorescence Images were taken after 48 hours with a 60x oil emulsion objective, Exposure time 1.25 seconds and EM gain of 675. Corresponding brightfield images were contrast enhanced with a phase filter (Ph3). Image processing and compositing was conducted in FIJI (Schindelin *et al*., 2012).

### Pellicle-to-pellicles assays

The separate pellicle pairs were placed within the same petri dish with 5 ml HS media and incubated at 30°C with gentle shaking of 60 RPM. After 24 hours the pellicle pairs were removed, and fluorescence imagery taken. The Boundary detection assay used a larger pellicle, grown for 2 days in a 10×10 cm square petri dish with 50 ml HS media. To produce a uniform pellicle at this larger surface area, a 1:10 ration of the liquid from under the first batch of pellicles was used as inoculum. Sender and receiver pellicles were removed from cultures, excess media was removed, and the pellicles were sliced in half along the diagonal. To facilitate diffusion yet prevent sinking, the two pellicles were placed in proximity on soft 0.6% HS agar and incubated at 30°C for 24 hours before fluorescence imagery was taken.

## Acknowledgements

The authors wish to thank Dr Benjamin Reeve and Dr Carlos Bricio-Garberi for initial help with this project and acknowledge the UK Engineering and Physcial Sciences Research Council (EPSRC) for the funding of this work.

## Author Contributions

V.J.G., K.T.W. and T.E. conceived and designed the experiments, K.T.W., A.D. and A.G. performed the experiments, K.T.W., V.J.G. and A.D. analysed the experimental data, V.J.G., K.T.W. and T.E. prepared the manuscript.

## References

Basu, S., Gerchman, Y., Collins, C.H., Arnold, F.H., Weiss, R., (2005) A synthetic multicellular system for programmed pattern formation. Nature 434: 1130–1134.

Boehm, C.R., Grant, P.K., Haseloff, J., (2018) Programmed hierarchical patterning of bacterial populations. Nat Commun 9: 776.

Brophy, J.A.N., Voigt, C.A., (2014) Principles of genetic circuit design. Nat Methods 11: 508–520.

Cameron, D.E., Bashor, C.J., Collins, J.J., (2014) A brief history of synthetic biology. Nat Rev Microbiol 12: 381–390.

Canton, B., Labno, A., Endy, D., (2008) Refinement and standardization of synthetic biological parts and devices. Nat Biotechnol 26: 787–793.

Chen, Y., Su, N., Zhang, K., Zhu, S., Zhao, L., Fang, F., Ren, L., Guo, Y., (2017) In-Depth Analysis of the Structure and Properties of Two Varieties of Natural Luffa Sponge Fibers. Materials (Basel) 10: 479.

Churchill, M.E.A., Sibhatu, H.M., Uhlson, C.L., (2011) Defining the structure and function of acyl-homoserine lactone autoinducers. Methods Mol Biol 692: 159–171.

Florea, M., Hagemann, H., Santosa, G., Abbott, J., Micklem, C.N., Spencer-Milnes, X., de Arroyo Garcia, L., Paschou, D., Lazenbatt, C., Kong, D., Chughtai, H., Jensen, K., Freemont, P.S., Kitney, R., Reeve, B., Ellis, T., (2016) Engineering control of bacterial cellulose production using a genetic toolkit and a new cellulose-producing strain. Proc Natl Acad Sci 113: E3431–E3440.

Jang, W.D., Hwang, J.H., Kim, H.U., Ryu, J.Y., Lee, S.Y., (2017) Bacterial cellulose as an example product for sustainable production and consumption. Microb Biotechnol 10: 1181–1185.

Karig, D., Martini, K.M., Lu, T., DeLateur, N.A., Goldenfeld, N., Weiss, R., (2018) Stochastic Turing patterns in a synthetic bacterial population. Proc Natl Acad Sci 115: 6572–6577.

Lee, K.Y., Buldum, G., Mantalaris, A., Bismarck, A., (2014) More Than Meets the Eye in Bacterial Cellulose: Biosynthesis, Bioprocessing, and Applications in Advanced Fiber Composites. Macromol Biosci 14: 10–32.

Liu, C., Fu, X., Liu, L., Ren, X., Chau, C.K.L., Li, S., Xiang, L., Zeng, H., Chen, G., Tang, L.H., Lenz, P., Cui, X., Huang, W., Hwa, T., Huang, J.D., (2011) Sequential establishment of stripe patterns in an expanding cell population. Science 334: 238–241.

Liu, M., Liu, L., Jia, S., Li, S., Zou, Y., Zhong, C., (2018) Complete genome analysis of Gluconacetobacter xylinus CGMCC 2955 for elucidating bacterial cellulose biosynthesis and metabolic regulation. Sci Rep 8: 6266.

Nguyen, P.Q., Botyanszki, Z., Tay, P.K.R., Joshi, N.S., (2014) Programmable biofilm-based materials from engineered curli nanofibres. Nat Commun 5: 4945.

Nguyen, P.Q., Courchesne, N.M.D., Duraj-Thatte, A., Praveschotinunt, P., Joshi, N.S., (2018) Engineered Living Materials: Prospects and Challenges for Using Biological Systems to Direct the Assembly of Smart Materials. Adv Mater 30: 1704847.

Park, J.P., Choi, M.J., Kim, S.H., Lee, S.H., Lee, H., (2014) Preparation of sticky Escherichia coli through surface display of an adhesive catecholamine moiety. Appl Environ Microbiol 80: 43–53.

Römling, U., Galperin, M.Y., (2015) Bacterial cellulose biosynthesis: diversity of operons, subunits, products, and functions. Trends Microbiol 23: 545–557.

Schindelin, J., Arganda-Carreras, I., Frise, E., Kaynig, V., Longair, M., Pietzsch, T., Preibisch, S., Rueden, C., Saalfeld, S., Schmid, B., Tinevez, J.Y., White, D.J., Hartenstein, V., Eliceiri, K., Tomancak, P., Cardona, A., (2012) Fiji: an open-source platform for biological-image analysis. Nat Methods 9: 676–682.

Scholes, N.S., Isalan, M., (2017) A three-step framework for programming pattern formation. Curr Opin Chem Biol 40: 1–7.

Scott, S.R., Hasty, J., (2016) Quorum Sensing Communication Modules for Microbial Consortia. ACS Synth Biol 5: 969–977.

Turing, A.M., (1952) The Chemical Basis of Morphogenesis. Philos Trans R Soc B Biol Sci 237: 37–72.

Waters, C.M., Bassler, B.L., (2005) Quorum sensing: cell-to-cell communication in bacteria. Annu Rev Cell Dev Biol 21: 319–346.

Wolpert, L., (1969) Positional information and the spatial pattern of cellular differentiation. J Theor Biol 25: 1–47.

Youssefian, S., Rahbar, N., (2015) Molecular Origin of Strength and Stiffness in Bamboo Fibrils. Sci Rep 5: 11116.

